# Habitat structure and social behavior - the development of social phenotypes in the desert locust

**DOI:** 10.1101/2025.09.27.677662

**Authors:** Madhansai Narisetty, Sercan Sayin, Yvonne Hertenberger, Stephen J. Simpson, Ahmed El Hady, Einat Couzin-Fuchs

## Abstract

Habitat structure has fundamental implications for behavior, foraging dynamics, and the social organization of animals. Locusts, which exhibit both a sedentary solitarious morph and a gregarious swarm-forming one, provide a powerful model to investigate how environmental factors shape social phenotypes. We conducted long-term continuous monitoring of locusts raised in semi-natural arenas with either clumped or distributed resources. Our findings reveal that clumped overnight roosting led to an increased propensity for coordinated marching the following day, a key precursor of swarm formation. These patterns became clearer during larval development, driving divergence in social affinity and group dynamics between habitat groups. At the molecular level, transcriptomic analysis identified a narrow set of differentially expressed genes, with enrichment in neuromodulatory and metabolic pathways, including candidates linked to monoaminergic signaling and juvenile hormone biology, such as a takeout-like JH-binding gene. These results demonstrate that spatial resource configuration is sufficient for lasting differences in individual and group dynamics and offer insight into the molecular processes mediating spontaneous, habitat-driven, phenotypic changes in a devastating plague insect.

## 1 Introduction

From global warming to land-use policy shifts, habitats are undergoing rapid transformations, profoundly impacting the survival and behavior of their inhabitants (Blowes et al., 2019; Clotfelter et al., 2004; Liu et al., 2024). Habitat structure influences how resident and migratory species navigate and utilize their environment, determines how populations are dispersed, and shapes the rate and nature of social interactions (He et al., 2019). Over longer timescales, environmentally mediated changes in interaction patterns and social structures impose selective pressures that drive phenotypic adaptations (Bailey and Moore, 2018; Testard et al., 2024). Advances in large-scale monitoring of animal movement and habitat characteristics now allow for clearer associations to be drawn between environmental features and behavioral patterns. However, while these techniques offer important insights into how factors such as topology, resource availability, and habitat heterogeneity are linked to individual and collective behaviors (Papageorgiou et al., 2021; Strandburg-Peshkin et al., 2017), the long-term behavioral, physiological, and socioecological consequences remain difficult to assess due to lack of continuous access to population-level behavioral data and limitations on field-omics.

Locusts exhibit strong adaptations to environmental fluctuations. Locust plagues are initiated when abnormal rainfall in arid environments creates favorable breeding conditions (Cressman and Stefanski, 2016). When resources become scarce, locusts crowd onto the remaining vegetation, triggering behavioral and physiological adaptations to high-density conditions (Ellis and Ashall, 1957; Roffey and Popov, 1968). This adaptation, known as phase polyphenism, refers to the transition from a low-density, sedentary, camouflaged solitarious phenotype to a highly active, conspicuous, swarm-forming gregarious phenotype in response to increased population density (Lester et al., 2005; Pener and Simpson, 2009; Simpson et al., 2005). It is now clear that this phenotypic plasticity is central to the formation of migratory locust swarms (Cullen et al., 2017; Pener and Simpson, 2009; Sword et al., 2000) with ample evidence that food availability, distribution, and quality are important factors (Bazazi et al., 2011; Despland et al., 2000; Despland and Simpson, 2000a). Consistent with field observations of gregarious aggregations in patchy environments, semi-field, laboratory, and modeling studies have shown that clumped resources enhance the expression of the gregarious state (Bouaichi et al., 1996; Collett et al., 1998; Despland et al., 2000; Despland and Simpson, 2000a). These studies highlighted how fine-scale habitat features can shape behavior, while also pointing to the influence of resource type (Bouaichi et al., 1996), food quality (Despland and Simpson, 2000b; Simpson, 2022), group size (Collett et al., 1998), and exposure time.

In addition to the humanitarian importance of locust research for improving outbreak prediction and control, locusts provide a tractable model system for studying social plasticity and its environmental drivers. While unusually rapid in locusts, adaptations in physiology and behavior to changing social conditions are widespread in the animal kingdom (Hall, 1998; Yadav et al., 2024). Comparative transcriptomics of solitarious and gregarious colonies has revealed widespread differential gene expression (Bakkali and Martín-Blázquez, 2018; Yang et al., 2019), including changes in neuromodulatory systems (such as serotonin, dopamine, glutamate and GABA (Anstey et al., 2009; Ma et al., 2011; Yang et al., 2023)) and endocrine pathways such as juvenile hormone and related targets (Guo et al., 2020; Wang and Kang, 2014). Yet these datasets typically compare categorical laboratory treatments (larger crowded cages versus small confined solitary containers). It is not yet known to what extent the same pathways regulate phenotypic changes under natural, environmentally mediated fluctuations in local density, nor how these link to movement and foraging behaviors. Furthermore, previous studies almost exclusively relied on one type of behavioral paradigm, which we propose to expand to additionally capture inter-individual interactions and group dynamics. To address this, we built on previous ideas and results (Despland et al., 2000) using current techniques for continuous population-level tracking, combined with group-level behavioral analysis and RNA sequencing in semi-natural arenas. Our results confirm previous findings and show that habitat structure has a rapid and long-lasting effect on locust group dynamics and gene expression, providing a framework to investigate the mechanisms underlying social phenotype differentiation in ecologically relevant settings.

## 2 Results

We investigated how habitat structure is linked to the development of locust phenotypes by raising locusts in observation arenas with clumped or distributed resource conditions (food and roosting sites, Fig. 1A). The arenas were designed for continuous behavioral monitoring throughout development, from the second to the final larval instar (n = 150 locusts per condition). Apart from resource distribution, which remained constant throughout the trials, all other conditions, including total food volume and total roosting areas, were standardized to be similar between treatments [i.e in Trial 1, 140 grams in each arena exchanged daily, 500 cm^2^ total roosting area]. This was reflected by similar mortality, development rate, and food consumption for the two conditions, monitored daily (Supplementary Figure 3).

**Figure 1:**
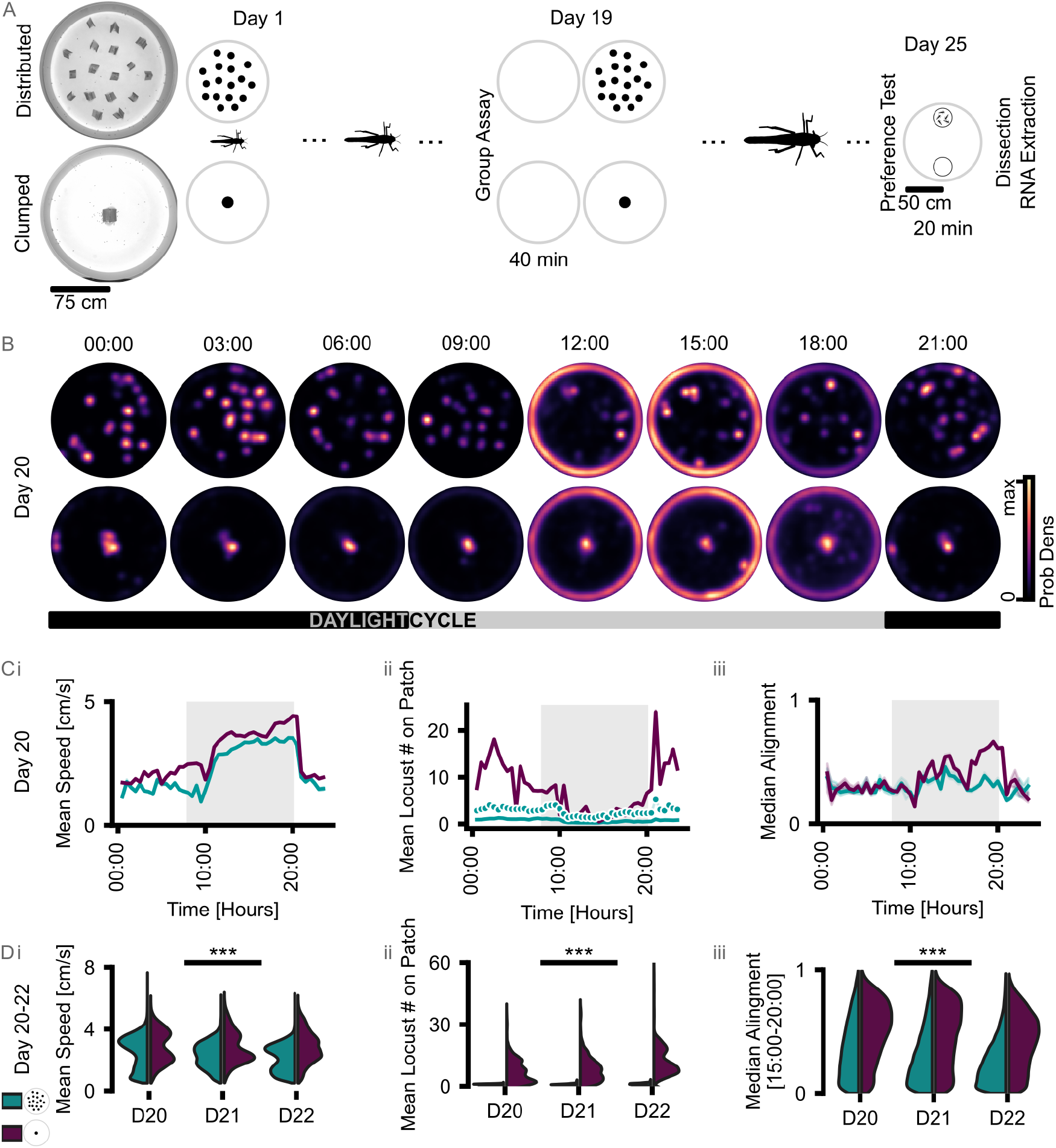
Experimental setup and population-level activity patterns. (A) Schematic of the experimental layout, the structure of resources in the central/clumped and distributed observation arenas, and experimental timeline. (B) Example density maps of locust positions over a 24-hour period (day 20) illustrate characteristic diurnal patterns. Animal density was estimated based on individual trajectories in 3 hour intervals extracted via the motion capture system. (C) Daily population metrics during the last week of the experiment (same day as in B). Time serious of (i) mean speed of moving individuals, (ii) number of animals on food/roosting patches, and (iii) an alignment index (scaled difference between aligned vs misaligned neighbor pairs) of all tracked locusts from the clumped (purple) and distributed (blue) condition. Shade light-on period. (D) Distributions of the same three metrics for three consecutive days (day 20-22). For statistics, linear mixed-effects models with random intercepts per experimental session were used to account for repeated measurements across days. Please see Table 1 for statistical summaries.

Example density maps over a 24-hour period illustrate characteristic diurnal activity patterns and locust distributions in both arena types (Fig. 1B, day 20 of the experiment). Strongly coupled to the light-dark cycle, locusts in both conditions spent the majority of the dark hours at the roosting patches and displayed increased activity, such as marching along arena walls and across the arenas, during daylight hours. These patterns remained generally similar over days (Fig. 1D, distributions for three consecutive days in the final week of the experiment in which locusts were marked with IR markers for continuous tracking including overnight; see also Fig. 1C for time series)). Locusts in the clumped resource condition exhibited higher levels of activity (mean speed) and higher degree of pairwise alignment among moving neighbors during the active hours (24-h traces in Fig. 1C; inter-day comparison in Fig. 1D; for speed and alignment, animals on patches and along the arena border were excluded).

**Table 1:**
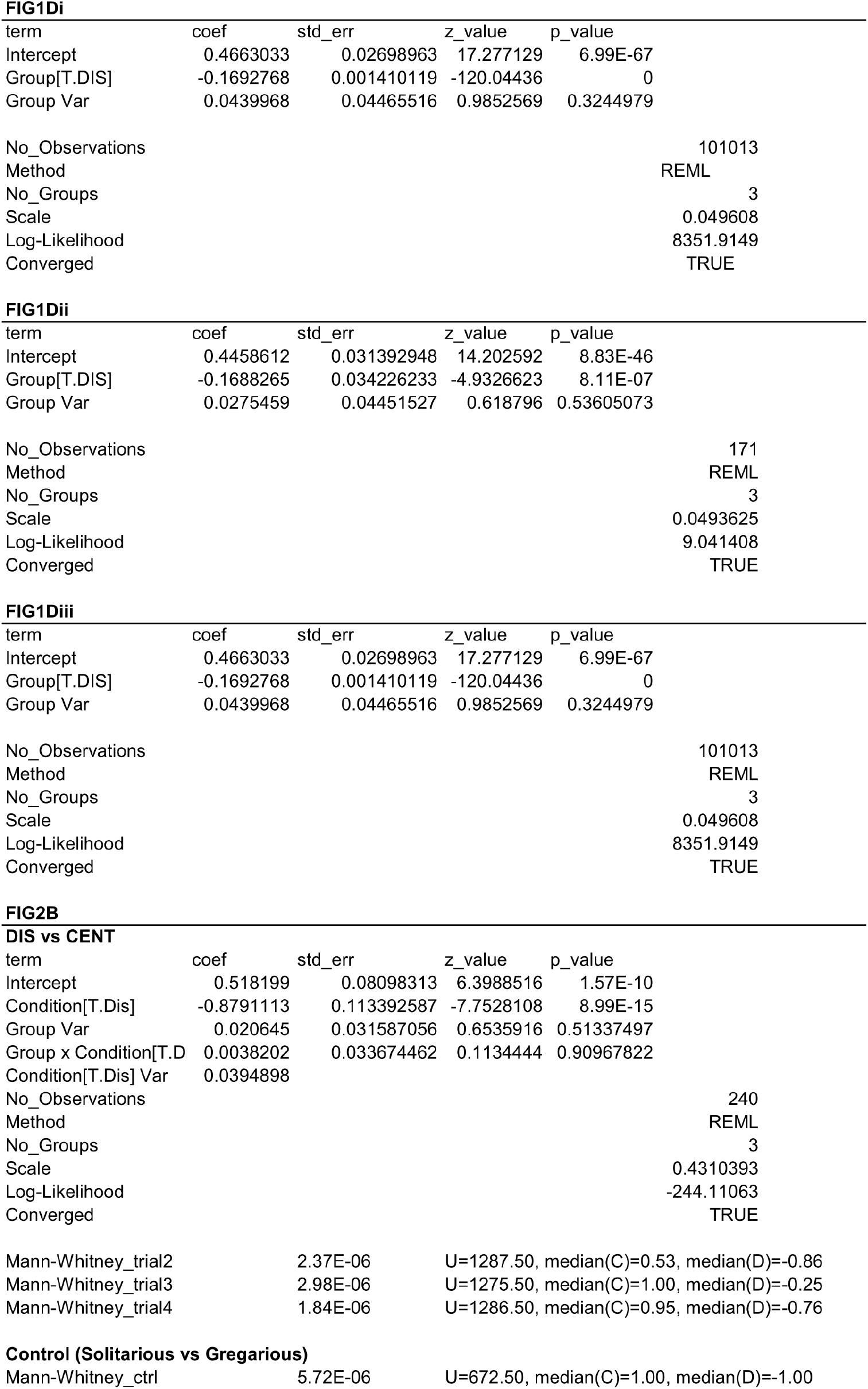
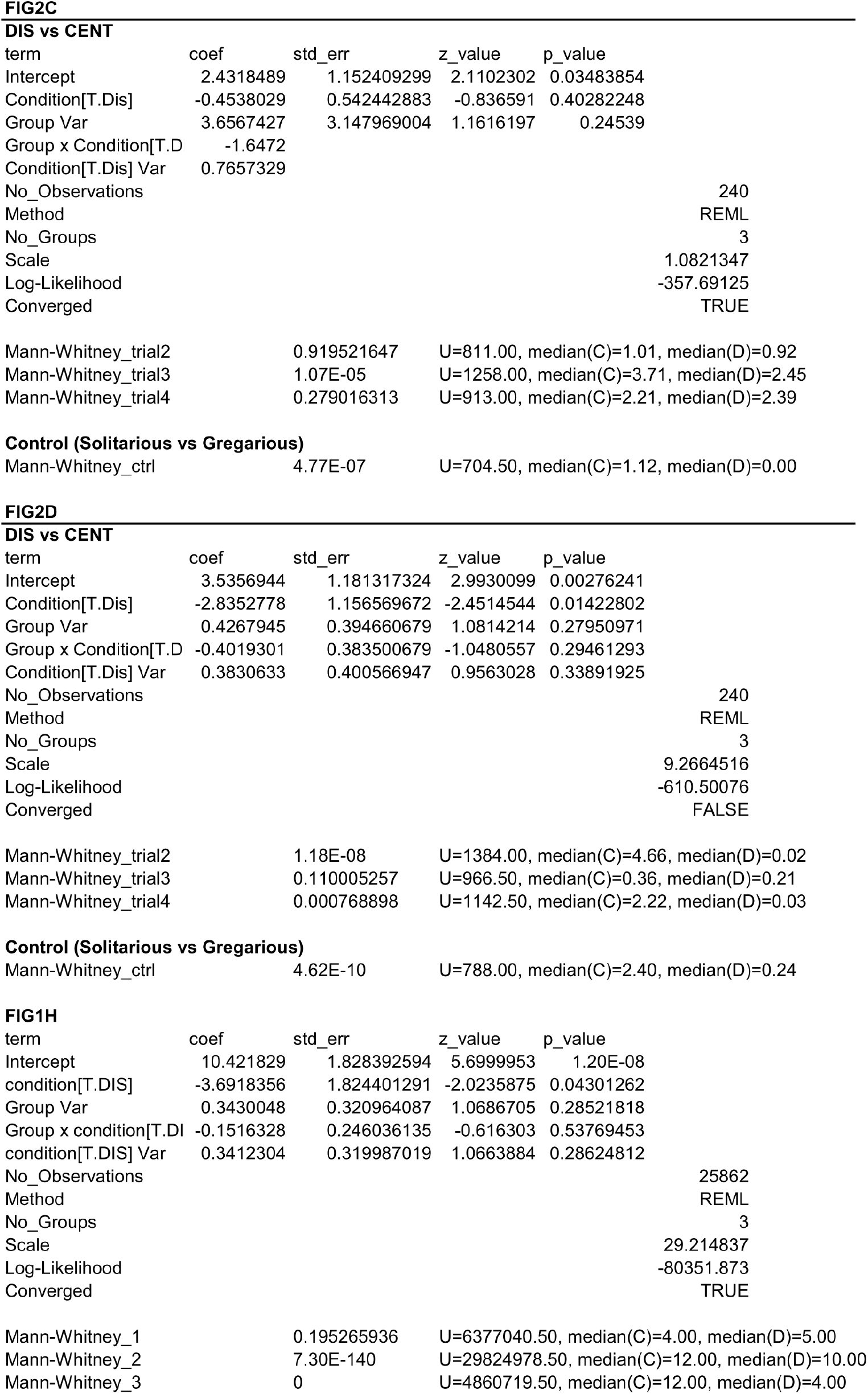
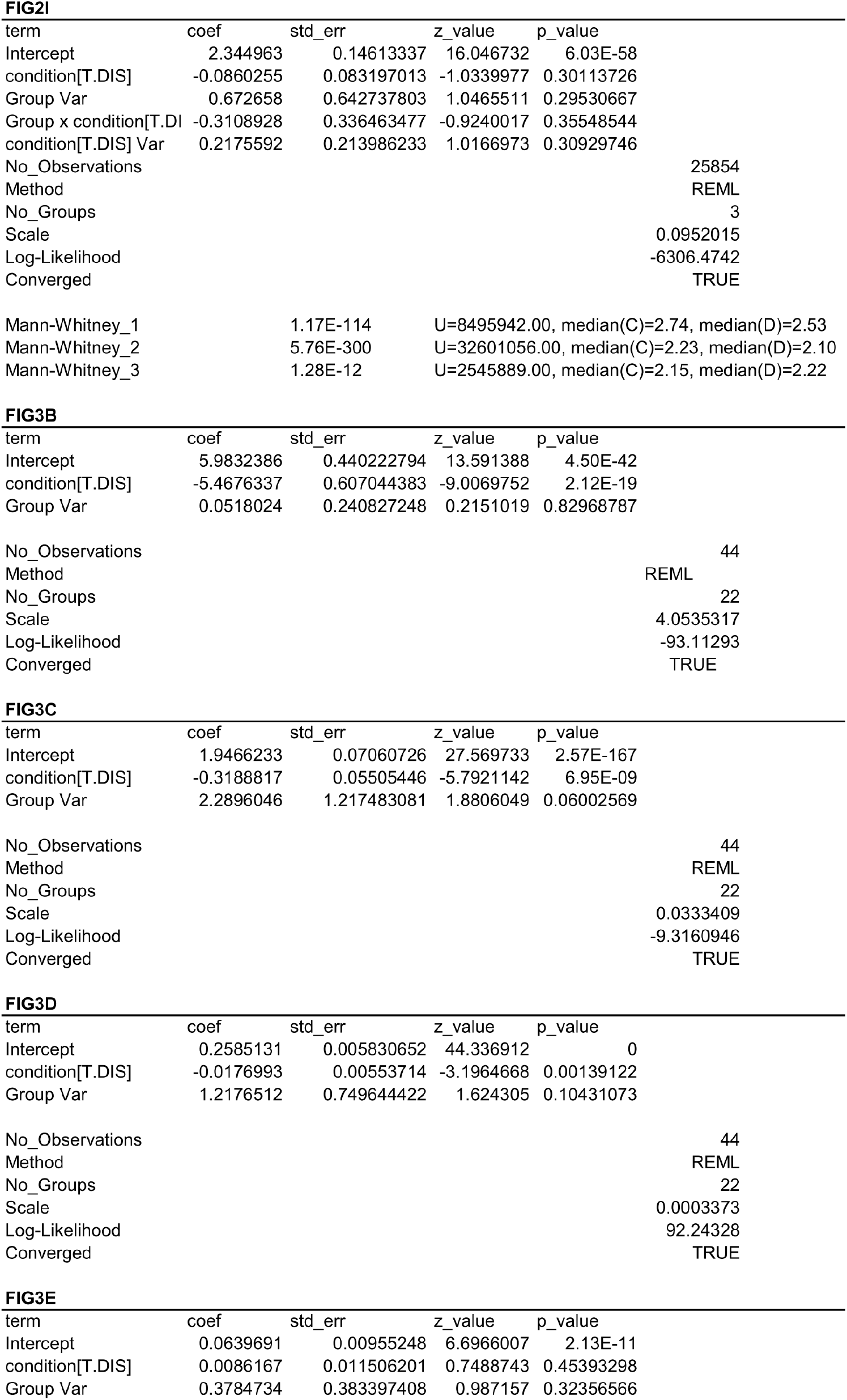
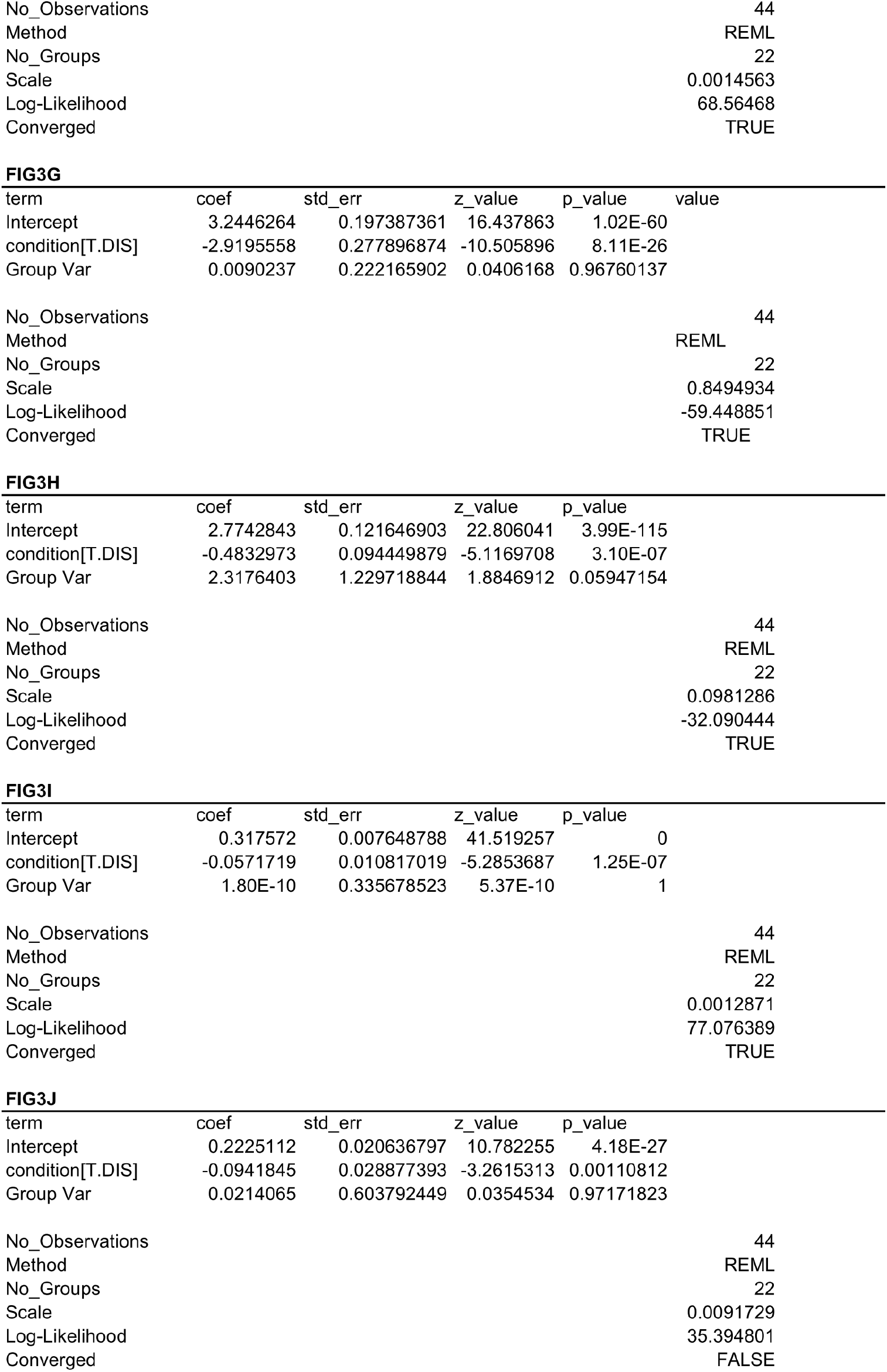
Statistical summaries.

**FIG1Di**
| term | coef | std_err | z_value | p_value |
| --- | --- | --- | --- | --- |
| Intercept | 0.4663033 | 0.02698963 | 17.277129 | 6.99E-67 |
| Group[T.DIS] | -0.1692768 | 0.001410119 | -120.04436 | 0 |
| Group Var | 0.0439968 | 0.04465516 | 0.9852569 | 0.3244979 |
| No_Observations |  |  |  | 101013 |
| Method |  |  |  | REML |
| No_Groups |  |  |  | 3 |
| Scale |  |  |  | 0.049608 |
| Log-Likelihood |  |  |  | 8351.9149 |
| Converged |  |  |  | TRUE |

**FIG1Dii**
| term | coef | std_err | z_value | p_value |
| --- | --- | --- | --- | --- |
| Intercept | 0.4458612 | 0.031392948 | 14.202592 | 8.83E-46 |
| Group[T.DIS] | -0.1688265 | 0.034226233 | -4.9326623 | 8.11E-07 |
| Group Var | 0.0275459 | 0.04451527 | 0.618796 | 0.53605073 |
| No_Observations |  |  |  | 171 |
| Method |  |  |  | REML |
| No_Groups |  |  |  | 3 |
| Scale |  |  |  | 0.0493625 |
| Log-Likelihood |  |  |  | 9.041408 |
| Converged |  |  |  | TRUE |

**FIG2B**
| **DIS vs CENT** |  |  |  |  |
| --- | --- | --- | --- | --- |
| term | coef | std_err | z_value | p_value |
| Intercept | 0.518199 | 0.08098313 | 6.3988516 | 1.57E-10 |
| Condition[T.Dis] | -0.8791113 | 0.113392587 | -7.7528108 | 8.99E-15 |
| Group Var | 0.020645 | 0.031587056 | 0.6535916 | 0.51337497 |
| Group x Condition[T.D | 0.0038202 | 0.033674462 | 0.1134444 | 0.90967822 |
| Condition[T.Dis] Var | 0.0394898 |  |  |  |
| No_Observations |  |  |  | 240 |
| Method |  |  |  | REML |
| No_Groups |  |  |  | 3 |
| Scale |  |  |  | 0.4310393 |
| Log-Likelihood |  |  |  | -244.11063 |
| Converged |  |  |  | TRUE |
| Mann-Whitney_trial2 |  | 2.37E-06 | U=1287.50, median(C)=0.53, median(D)=-0.86 |  |
| Mann-Whitney_trial3 |  | 2.98E-06 | U=1275.50, median(C)=1.00, median(D)=-0.25 |  |
| Mann-Whitney_trial4 |  | 1.84E-06 | U=1286.50, median(C)=0.95, median(D)=-0.76 |  |

: Table 1: Statistical summaries.
|  |  |  |
| --- | --- | --- |
| Mann-Whitney_ctrl | 5.72E-06 | U=672.50, median(C)=1.00, median(D)=-1.00 |
| Intercept | 2.4318489 | 1.152409299 | 2.1102302 | 0.03483854 |
| Condition[T.Dis] | -0.4538029 | 0.542442883 | -0.836591 | 0.40282248 |
| Group Var | 3.6567427 | 3.147969004 | 1.1616197 | 0.24539 |
| Group x Condition[T.D | -1.6472 |  |  |  |
| Condition[T.Dis] Var | 0.7657329 |  |  |  |
| No_Observations |  |  |  | 240 |
| Method |  |  |  | REML |
| No_Groups |  |  |  | 3 |
| Scale |  |  |  | 1.0821347 |
| Log-Likelihood |  |  |  | -357.69125 |
| Converged |  |  |  | TRUE |
| Mann-Whitney_trial2 |  | 0.919521647 | U=811.00, median(C)=1.01, median(D)=0.92 |  |
| Mann-Whitney_trial3 |  | 1.07E-05 | U=1258.00, median(C)=3.71, median(D)=2.45 |  |
| Mann-Whitney_trial4 |  | 0.279016313 | U=913.00, median(C)=2.21, median(D)=2.39 |  |

**Control (Solitarious vs Gregarious)**
|  |  |  |
| --- | --- | --- |
| Mann-Whitney_ctrl | 4.77E-07 | U=704.50, median(C)=1.12, median(D)=0.00 |

| term | coef | std_err | z_value | p_value |
| --- | --- | --- | --- | --- |
| Intercept | 3.5356944 | 1.181317324 | 2.9930099 | 0.00276241 |
| Condition[T.Dis] | -2.8352778 | 1.156569672 | -2.4514544 | 0.01422802 |
| Group Var | 0.4267945 | 0.394660679 | 1.0814214 | 0.27950971 |
| Group x Condition[T.D | -0.4019301 | 0.383500679 | -1.0480557 | 0.29461293 |
| Condition[T.Dis] Var | 0.3830633 | 0.400566947 | 0.9563028 | 0.33891925 |
| No_Observations |  |  |  | 240 |
| Method |  |  |  | REML |
| No_Groups |  |  |  | 3 |
| Scale |  |  |  | 9.2664516 |
| Log-Likelihood |  |  |  | -610.50076 |
| Converged |  |  |  | FALSE |
| Mann-Whitney_trial2 |  | 1.18E-08 | U=1384.00, median(C)=4.66, median(D)=0.02 |  |
| Mann-Whitney_trial3 |  | 0.110005257 | U=966.50, median(C)=0.36, median(D)=0.21 |  |
| Mann-Whitney_trial4 |  | 0.000768898 | U=1142.50, median(C)=2.22, median(D)=0.03 |  |

**Control (Solitarious vs Gregarious)**
|  |  |  |
| --- | --- | --- |
| Mann-Whitney_ctrl | 4.62E-10 | U=788.00, median(C)=2.40, median(D)=0.24 |

: Table 1: Statistical summaries continued.
| term | coef | std_err | z_value | p_value |
| --- | --- | --- | --- | --- |
| Intercept | 10.421829 | 1.828392594 | 5.6999953 | 1.20E-08 |
| condition[T.DIS] | -3.6918356 | 1.824401291 | -2.0235875 | 0.04301262 |
| Group Var | 0.3430048 | 0.320964087 | 1.0686705 | 0.28521818 |
| Group x condition[T.DI | -0.1516328 | 0.246036135 | -0.616303 | 0.53769453 |
| condition[T.DIS] Var | 0.3412304 | 0.319987019 | 1.0663884 | 0.28624812 |
| No_Observations |  |  |  | 25862 |
| Method |  |  |  | REML |
| No_Groups |  |  |  | 3 |
| Scale |  |  |  | 29.214837 |
| Log-Likelihood |  |  |  | -80351.873 |
| Converged |  |  |  | TRUE |
| Mann-Whitney_1 |  | 0.195265936 | U=6377040.50, median(C)=4.00, median(D)=5.00 |  |
| Mann-Whitney_2 |  | 7.30E-140 | U=29824978.50, median(C)=12.00, median(D)=10.00 |  |
| Mann-Whitney_3 |  | 0 | U=4860719.50, median(C)=12.00, median(D)=4.00 |  |

**FIG2I**
| term | coef | std_err | z_value | p_value |
| --- | --- | --- | --- | --- |
| Intercept | 2.344963 | 0.14613337 | 16.046732 | 6.03E-58 |
| condition[T.DIS] | -0.0860255 | 0.083197013 | -1.0339977 | 0.30113726 |
| Group Var | 0.672658 | 0.642737803 | 1.0465511 | 0.29530667 |
| Group x condition[T.DI | -0.3108928 | 0.336463477 | -0.9240017 | 0.35548544 |
| condition[T.DIS] Var | 0.2175592 | 0.213986233 | 1.0166973 | 0.30929746 |
| No_Observations |  |  |  | 25854 |
| Method |  |  |  | REML |
| No_Groups |  |  |  | 3 |
| Scale |  |  |  | 0.0952015 |
| Log-Likelihood |  |  |  | -6306.4742 |
| Converged |  |  |  | TRUE |
| Mann-Whitney_1 |  | 1.17E-114 | U=8495942.00, median(C)=2.74, median(D)=2.53 |  |
| Mann-Whitney_2 |  | 5.76E-300 | U=32601056.00, median(C)=2.23, median(D)=2.10 |  |
| Mann-Whitney_3 |  | 1.28E-12 | U=2545889.00, median(C)=2.15, median(D)=2.22 |  |

**FIG3B**
| term | coef | std_err | z_value | p_value |
| --- | --- | --- | --- | --- |
| Intercept | 5.9832386 | 0.440222794 | 13.591388 | 4.50E-42 |
| condition[T.DIS] | -5.4676337 | 0.607044383 | -9.0069752 | 2.12E-19 |
| Group Var | 0.0518024 | 0.240827248 | 0.2151019 | 0.82968787 |
| No_Observations |  |  |  | 44 |
| Method |  |  |  | REML |
| No_Groups |  |  |  | 22 |
| Scale |  |  |  | 4.0535317 |
| Log-Likelihood |  |  |  | -93.11293 |
| Converged |  |  |  | TRUE |

**FIG3C**
| term | coef | std_err | z_value | p_value |
| --- | --- | --- | --- | --- |
| Intercept | 1.9466233 | 0.07060726 | 27.569733 | 2.57E-167 |
| condition[T.DIS] | -0.3188817 | 0.05505446 | -5.7921142 | 6.95E-09 |
| Group Var | 2.2896046 | 1.217483081 | 1.8806049 | 0.06002569 |
| No_Observations |  |  |  | 44 |
| Method |  |  |  | REML |
| No_Groups |  |  |  | 22 |
| Scale |  |  |  | 0.0333409 |
| Log-Likelihood |  |  |  | -9.3160946 |
| Converged |  |  |  | TRUE |

**FIG3D**
| term | coef | std_err | z_value | p_value |
| --- | --- | --- | --- | --- |
| Intercept | 0.2585131 | 0.005830652 | 44.336912 | 0 |
| condition[T.DIS] | -0.0176993 | 0.00553714 | -3.1964668 | 0.00139122 |
| Group Var | 1.2176512 | 0.749644422 | 1.624305 | 0.10431073 |
| No_Observations |  |  |  | 44 |
| Method |  |  |  | REML |
| No_Groups |  |  |  | 22 |
| Scale |  |  |  | 0.0003373 |
| Log-Likelihood |  |  |  | 92.24328 |
| Converged |  |  |  | TRUE |

: Table 1: Statistical summaries continued.
| term | coef | std_err | z_value | p_value |
| --- | --- | --- | --- | --- |
| Intercept | 0.0639691 | 0.00955248 | 6.6966007 | 2.13E-11 |
| condition[T.DIS] | 0.0086167 | 0.011506201 | 0.7488743 | 0.45393298 |
| Group Var | 0.3784734 | 0.383397408 | 0.987157 | 0.32356566 |

|  |  |
| --- | --- |
| No_Observations | 44 |
| Method | REML |
| No_Groups | 22 |
| Scale | 0.0014563 |
| Log-Likelihood | 68.56468 |
| Converged | TRUE |

FIG3G
| term | coef | std_err | z_value | p_value | value |
| --- | --- | --- | --- | --- | --- |
| Intercept | 3.2446264 | 0.197387361 | 16.437863 | 1.02E-60 |  |
| condition[T.DIS] | -2.9195558 | 0.277896874 | -10.505896 | 8.11E-26 |  |
| Group Var | 0.0090237 | 0.222165902 | 0.0406168 | 0.96760137 |  |

|  |  |
| --- | --- |
| No_Observations | 44 |
| Method | REML |
| No_Groups | 22 |
| Scale | 0.8494934 |
| Log-Likelihood | -59.448851 |
| Converged | TRUE |

FIG3H
| term | coef | std_err | z_value | p_value |
| --- | --- | --- | --- | --- |
| Intercept | 2.7742843 | 0.121646903 | 22.806041 | 3.99E-115 |
| condition[T.DIS] | -0.4832973 | 0.094449879 | -5.1169708 | 3.10E-07 |
| Group Var | 2.3176403 | 1.229718844 | 1.8846912 | 0.05947154 |

|  |  |
| --- | --- |
| No_Observations | 44 |
| Method | REML |
| No_Groups | 22 |
| Scale | 0.0981286 |
| Log-Likelihood | -32.090444 |
| Converged | TRUE |

FIG3I
| term | coef | std_err | z_value | p_value |
| --- | --- | --- | --- | --- |
| Intercept | 0.317572 | 0.007648788 | 41.519257 | 0 |
| condition[T.DIS] | -0.0571719 | 0.010817019 | -5.2853687 | 1.25E-07 |
| Group Var | 1.80E-10 | 0.335678523 | 5.37E-10 | 1 |

|  |  |
| --- | --- |
| No_Observations | 44 |
| Method | REML |
| No_Groups | 22 |
| Scale | 0.0012871 |
| Log-Likelihood | 77.076389 |
| Converged | TRUE |

FIG3J
| term | coef | std_err | z_value | p_value |
| --- | --- | --- | --- | --- |
| Intercept | 0.2225112 | 0.020636797 | 10.782255 | 4.18E-27 |
| condition[T.DIS] | -0.0941845 | 0.028877393 | -3.2615313 | 0.00110812 |
| Group Var | 0.0214065 | 0.603792449 | 0.0354534 | 0.97171823 |

: Table 1: Statistical summaries continued.
|  |  |
| --- | --- |
| No_Observations | 44 |
| Method | REML |
| No_Groups | 22 |
| Scale | 0.0091729 |
| Log-Likelihood | 35.394801 |
| Converged | FALSE |

To optimize visibility for tracking the locusts on the food patches, the first round of experiments was carried out using artificial diet patches laid flat in Petri dishes (T1, artificial/flat patches). The composition of the artificial diet—gelatin, lentil paste, honey, water, and tamarind paste— was selected based on preliminary assessments of survival and vitality. To account for potential diet-specific effects, the experiments were repeated twice more using natural wheatgrass sods (T2, fully grown wheatgrass in soil) and cut grass plates (T3, natural/flat patches). Developmental rates in all three trials were similar, allowing behavioral tests conducted at the end of each trial to be performed at similar developmental time points.

### 2.1 Habitat structure shapes individual and group behavior

Behavior was characterized at the end of the observation period using a two-choice preference assay, an adaptation of the Roessingh assay (Roessingh et al., 1993a), which has been widely used to describe locust phenotypes. We also used a group assay to capture inter-individual interactions and group-level behavior. Calculating an estimate of social affinity in the individual assay (Fig. 2A) based on the proportion of time a locust spent at the stimulus patch (containing eight gregarious nymphs) versus the empty side revealed a significantly stronger attraction for locusts raised in the clumped condition compared to those from the distributed condition (positive preference index; Fig. 2B, see also time spent at the stimulus patch in Fig. 2C and average heatmaps, Fig. 2A). These values were indistinguishable from those of control gregarious and solitarious locusts, respectively (i.e., locusts raised in high crowding and full isolation; Fig. 2B right, gray and black violins), reinforcing the concept of two distinct phenotypes over a continuous behavioral range. Nevertheless, unlike control gregarious and solitarious locusts, differences in non-social traits such as walking speed—a dominant phenotypic marker in locust literature—were minimal between the two habitat conditions (Fig. 2D).

**Figure 2:**
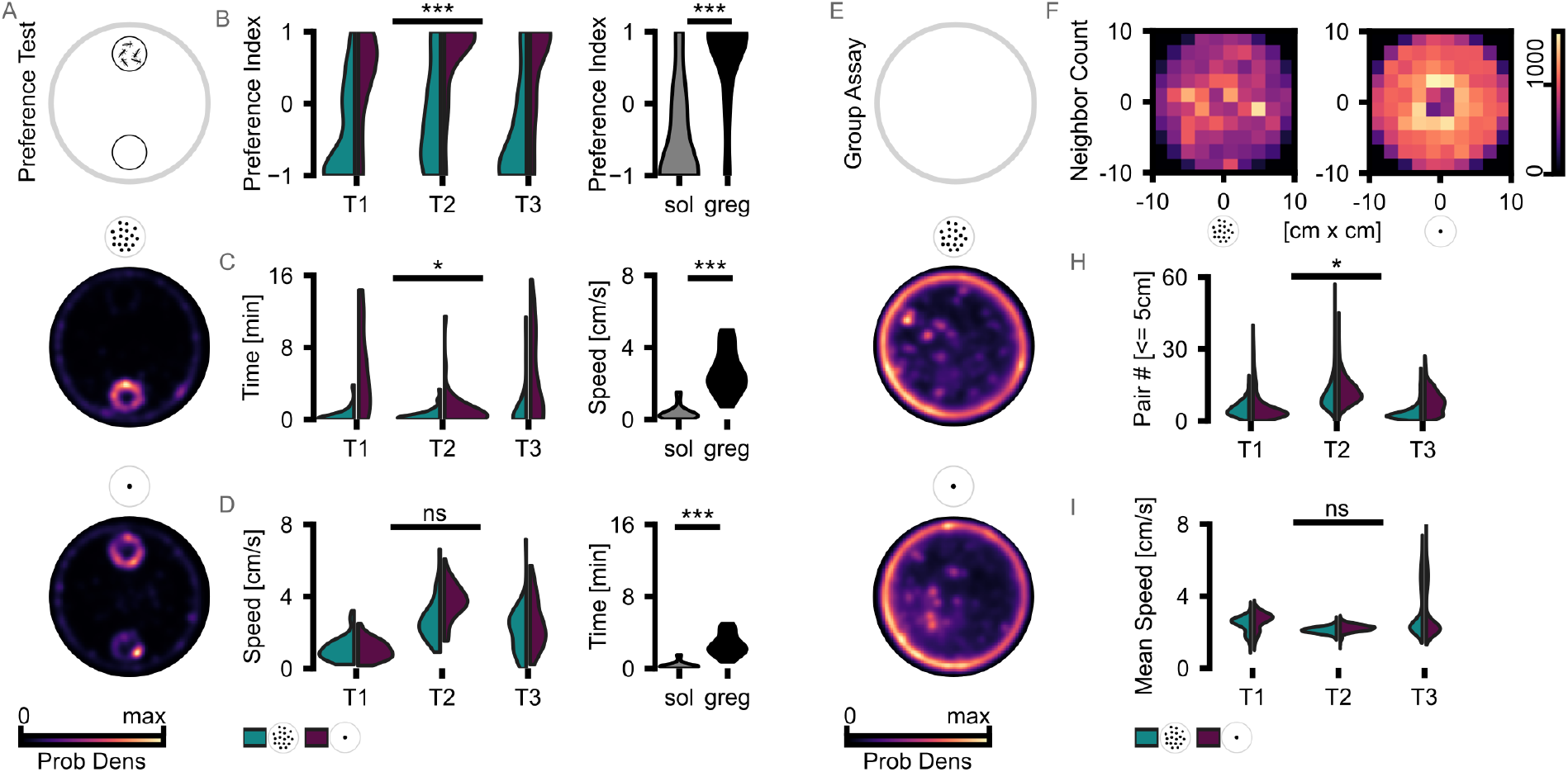
Behavioral phenotypes at the end of the experiment (A-D) Individual two-choice assay. Locusts were tested in a circular arena with two patches: one containing a group of gregarious nymphs and one empty; patch positions alternate across trials. (A) Schematics and averaged density heatmaps of all tested animals from the distributed and clumped arenas of trial 1 are plotted below. (B) Preference Index (PI) quantifying time near a conspecific stimulus (group of gregarious nymphs) vs an empty patch of all tested animals. Violin plots with PI distributions for the two conditions (distributed blue, clumped purple) in all three trials. Below are (C) the absolute time near the stimulus patch and (D) the average walking speed in all three sets of experiments (T1-T3). (B’-D’)respective values from locusts raised under typical solitarious and gregarious rearing conditions.(E-I) Group assay. (E) Example heatmaps of individuals in the empty arenas (F) local neighbor-density maps around individuals. (H) near-neighbor counts within 1 body length radius and mean walking speed (I). Linear mixed-effects models with random slopes and intercepts per trial were used to account for variability across independent experimental replicates. For comparing solitarious and gregarious control locust behavior, Mann-Whitney U tests were employed. Please see Table 1 for statistical summaries.

When tested in a group setting (open, empty arenas after removing all roosting sites), locusts from both conditions explored the arena with frequent interactions among individuals (Fig. 2E). Locusts from the clumped resource condition exhibited more frequent close-range interactions, as reflected in a higher local density around focal individuals (Fig. 2C, density maps averaged across all individuals in T1). Moreover, higher tendency to aggregate was observed across all trials (Fig 2E, total neighbor counts within a body length radius were significantly greater for locusts from the clumped condition (Fig. 2D, statistics in Table 1; linear mixed-effects models with random effects per trial)).

### 2.2 Behavioral patterns are reinforced along development

We then continued to study how these behavioral differences develop over the course of the experiment. As all three trials show similar behavioral metrics in both the preference test and the group assay, we focused our detailed analysis on the first trial, in which the flat artificial food patches allowed the best conditions for continuous population tracking. Differences in movement patterns between the arenas were already apparent early on, as expected given the different habitat structures; nevertheless, these became more pronounced as development progressed. We focused our analysis on two daily time windows: early morning, as locusts start leaving the patches (9:00–9:40), and late afternoon, when activity peaked (17:00–17:40). The 40-minute time slots were chosen to match the duration used for the group assays above. Average heatmaps for each observation week (Fig. 3A 3F) and daily population averages of the number of animals at the patches (Fig. 3B and G, dotted green shows the sum across all patches in the distributed condition), the overall speed (Fig. 3C and H), and the local alignment between nearest neighbors (Fig. 3D-E and I-J; scaled difference between aligned and misaligned pairs; see legend/methods for details) show that behavioral differences were apparent from the first days of the experiment and reinforced over time. We also observed strong day-to-day fluctuations, especially in the clumped resource condition, which appeared to coincide with developmental time points of synchronized molting events. Nevertheless, consistent differences in marching alignment were observed throughout the experiment, particularly during the afternoon active phase. Histograms of pairwise alignment scores (excluding wall-followers), color-coded by day, further illustrate this: in the clumped condition, the probability density shifts toward higher alignment values, indicating a higher likelihood of coordinated marching events as development progressed (Fig. 3J).

**Figure 3:**
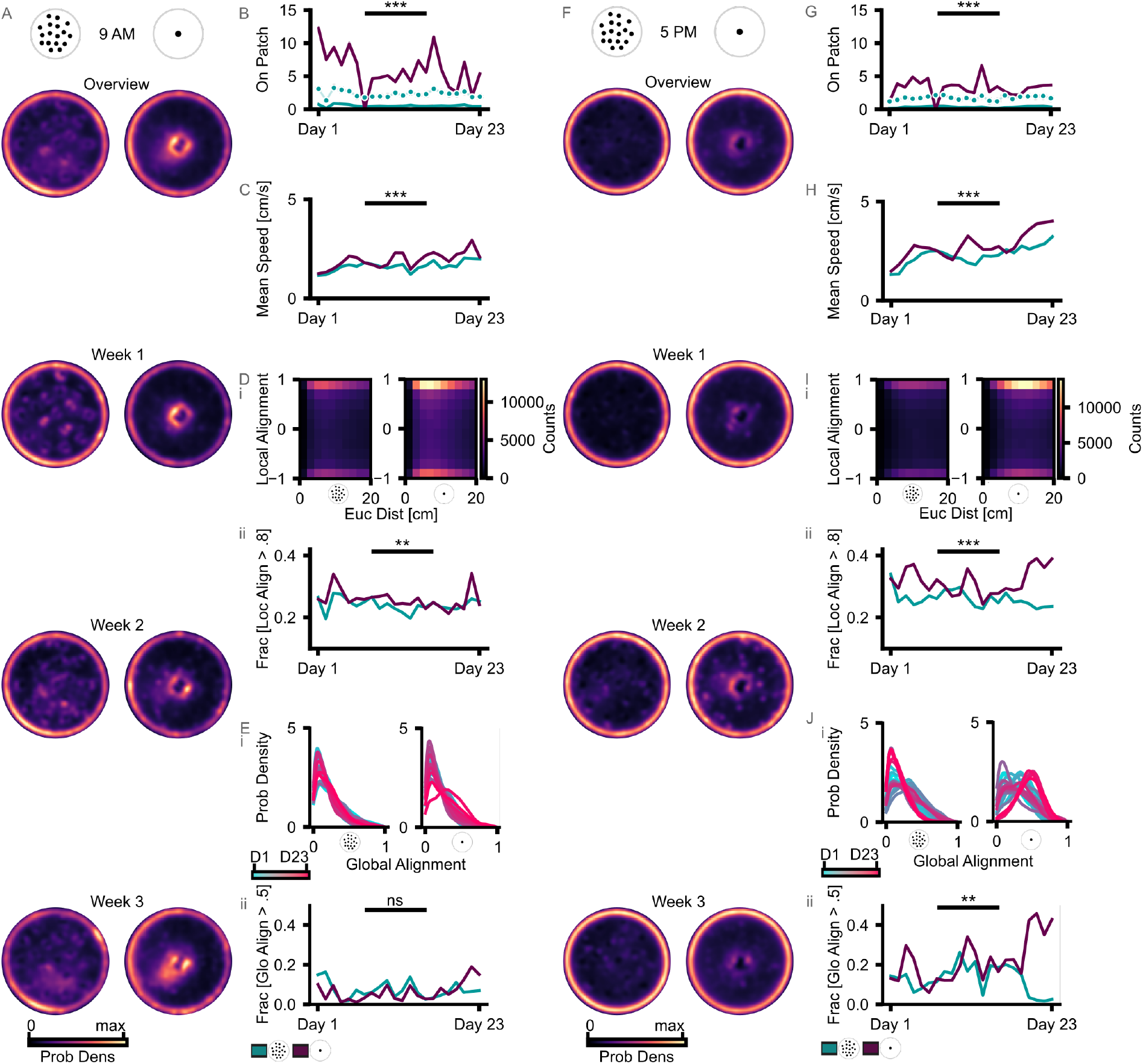
Development of behavioral phenotypes. Phenotypic differences manifest early on, yet differences accelerated on the third week. (A, F) Example density maps of locust positions over 23 days. Animal density was calculated based on individual trajectories in 40-minute chunks in a single day. Subpanels A-E display locust behavior at 9 am whereas F-J show at 5 pm. (B, G) Locust distributions across patch(es). (C, H) Hierarchical mean speeds across days. (D, I) Instantaneous local alignment between nearest neighbours. i: Local alignment has been calculated between 1 - dot product of two movement vectors and plotted here against Euclidean distance at the first frame. +1 denotes congruent motion, while -1 depicts opposite direction of movement. ii: Fraction of high alignment (> 0.8) across days. (E, J) Global alignment, calculated as in Fig. 1. Probability densities (i) and fraction of higher alignment (ii) for each day. Overall, for statistics, linear mixed-effects models with random intercepts per experimental session were used to account for repeated measurements across days. Please see Table 1for statistical summaries.

### 2.3 Transcriptomic profiling identifies candidate genes for the behavioral differences

To identify genes associated with the observed behavioral differences, we performed RNA sequencing on brain tissue collected at the end of the experiment. Differential expression analysis revealed 157 genes (log_2_ FoldChange > 0.5, adjusted *p*-value <10^−1^) that distinguish the two habitat conditions (Fig 4B). While we didn’t observe drastic differences in activity level in this study (i.e. speed differences were modest even when they were statistically significant), we nevertheless observed a number of DEGs involved in metabolism. Several genes in the DEG list were related to protein synthesis and development (Fig 4C), suggesting significant rewiring due to habitat-induced experience. Perhaps more significantly, neuronal activity is directly modulated by altering secondary messengers such as Rap guanine nucleotide exchange factor 4 (Fernandes et al., 2015), excitability (KCNQ-like potassium channels (Tsuboi et al., 2022; Wei et al., 1996)), TRPM channels (Hu and Wolfner, 2019; Turner et al., 2016), Cdk5R1 activator (Hahn et al., 2005), synaptic transmission (acetylcholinesterase (Wolfgang and Forte, 1989)), stoned-B (Fergestad et al., 1999), and futsch (Roos et al., 2000), providing a direct route for behavioral regulation.

**Figure 4:**
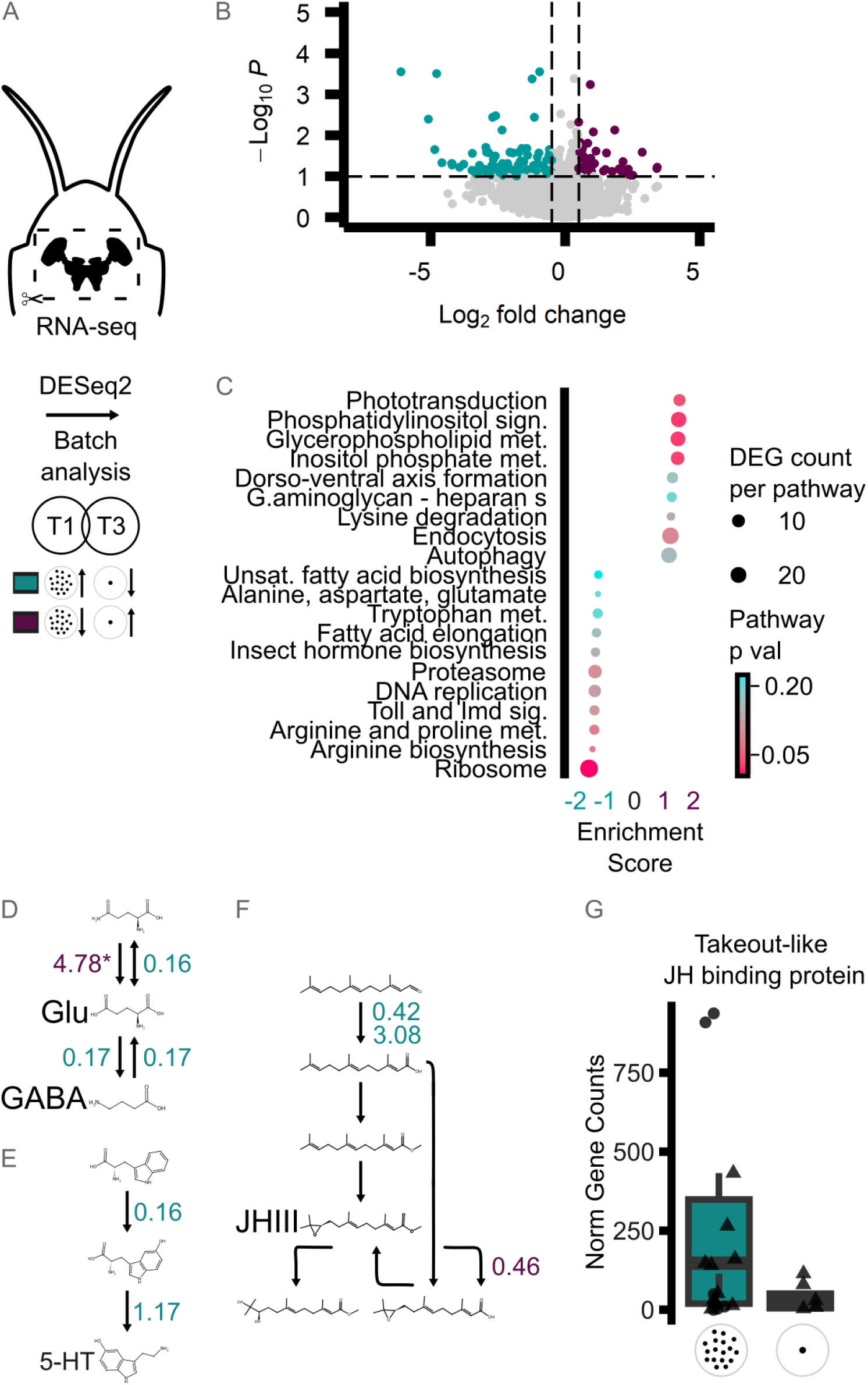
Transcriptional responses to habitat variation. (A) In the first and third trials, following the long-term monitoring of animal behavior, locust brains were dissected rapidly and the extracted RNAs were sequenced. Cyan: upregulated in distributed, purple: upregulated in centralized. (B) Differential gene expression analysis. Considering batch effects, DESeq2 revealed 99 genes upregulated in distributed and 58 genes in centralized conditions (log_2_ FoldChange > 0.5, adjusted *p*-value <10^−1^). (C) Gene set enrichment analysis (GSEA). The top 20 pathways are depicted here. (D–F) Mapping of DEGs (*p*-value < 0.1) onto individual enriched pathways. Values here show expression change at log_2_ scale. The asterisk marks the DEG with low brain-wide expression (*p* < 10^−5^). Juvenile hormone III expression is expected to be higher in distributed habitat due to higher precursor availability and lower expression of JHIII catalyzzing enzyme. (G) Juvenile hormone target Takeout-like protein (adjusted *p*-value <0.084) expression. Each point represents an individual sample within its respective trial.

Hypothesizing that neuromodulation is likely to take part in phase polyphenism, especially during the early stages in the transition between phases (Rogers et al., 2004), we investigated the top 20 pathways from the gene set enrichment analysis. We found trends in KEGG pathways for ‘alanine, aspartate, glutamate metabolism’, ‘tryptophan metabolism’, ‘insect hormone synthesis’; all enriched in the distributed condition (Fig 4B). Since KEGG pathways are extensive and include branches that do not directly contribute to a given neuromodulator’s synthesis or turnover, we mapped our DEGs with a relaxed threshold (*p*-value <10^−1^) to identify candidate targets. Despite enrichment at alanine/aspartate/glutamate metabolism, GABA-glutamate balance has not shifted (Fig 4D). The only major DEGs, a *glutaminase* (LOC1263358789, log_2_ FoldChange = −4.78), however it was expressed at a very low titer (baseMean = 10.5, marginally above the common threshold of 10 units). In ‘tryptophan metabolism’, two precursor enyzmes for serotonin were enriched at the distributed condition (Fig 4E). A stronger impact of habitat on gene expression was observed for Juvenile Hormone (JH) biosynthesis (Fig 4F). The strong change in the JH metabolitic pathway predicts higher JHIII level in the distributed condition which matches higher JH titers in solitarious individuals in both *L. migratoria* and *S. gregaria* (Tawfik et al., 2000; Wang and Kang, 2014; Wiesel et al., 1996). Takeout-like genes encoding JH binding proteins have been previously implicated in regulating behavioral plasticity and feeding in insects (Guo et al., 2011; Meunier et al., 2007) were found to be strongly upregulated in our distributed condition (Figs. 4F–G, Figure S2), pointing to selective endocrine (JH–takeout) plasticity as a major component of experience-dependent modulation.

## 3 Discussion

Although locust phenotypic plasticity reflects an adaptation to environmental change, its expression is fundamentally shaped by social interactions and collective dynamics. Understanding this process, therefore, requires studying behavior in naturalistic social environments across time. In this study, we examined the links between habitat structure, behavior, and the development of locust phenotypes, using semi-natural arenas that allow aggregations to spontaneously form and disperse. Our findings confirm the robustness of two distinct locust phenotypes, revisiting previous foundational work demonstrating the role of local habitat quality and structure on behavioural phase state both in the laboratory and the field (Bouaichi et al., 1996; Collett et al., 1998; Despland et al., 2000; Despland and Simpson, 2000a) with new tools that refine our understanding of their formation. We demonstrate that spatial habitat structure alone is sufficient to induce distinct, lasting behavioral phenotypes—in both individual behavior and group-level dynamics— that share similarities with lab-reared gregarious and solitarious colonies (Pener and Simpson, 2009; Sayin et al., 2025), yet develop under naturalistic density fluctuations and continuous social exposure. Social plasticity is widespread across the animal kingdom. Locusts provide a powerful model for investigating its underlying processes—a topic of growing interest in both neurobiology and ecology (Antunes et al., 2021; Fone and Porkess, 2008; Lihoreau et al., 2009; Yadav et al., 2024). For example, in the fruit fly, crowding induces lasting effects on life history and behavior linked to metabolic and developmental gene expression (Morimoto et al., 2023; Shipilina et al., 2024), while isolation impacts sleep and feeding through altered neuropeptide and biogenic amine signaling (Li et al., 2021; Yadav et al., 2024). In locusts, the behavioral effects of crowding and isolation have been functionally linked to regulation of neuromodulators and transmitters including dopamine (Guo et al., 2015; Ma et al., 2011; Yang et al., 2014) *in L*.*migratoria*, octopamine (Verlinden et al., 2010) and (Ma et al., 2015) in both locust species, glutamate-GABA (Yang et al., 2023) in *L*.*migratoria* and serotonin (Anstey et al., 2009; Simpson, 2022) in *S*.*gregaria*. However, these processes have mostly been studied under conditions of extreme isolation or crowding, and whether they similarly regulate behavioral plasticity under natural density fluctuations remains largely unexplored. Our observational approach addressed this by designing habitat configurations that better resemble real-life conditions. The fact that even under these moderate conditions—where locusts in both habitat types were exposed to and interacted with conspecifics—individuals segregated into two distinct phenotypes highlights the robustness of phase traits. Notably, since differences in local density were most pronounced during roosting and feeding periods, it is likely that these periods are particularly influential for social affinity traits. Tactile interactions at roosting sites can promote gregarious phenotypes via the previously described serotonin-initiated kinase–transcription cascade that affects gene expression and behavior (Anstey et al., 2009; Ott et al., 2012). Their coincidence with appetitive contexts (e.g., food) may additionally provide positive reinforcement, conditioning such interactions as rewarding. These non-mutually exclusive processes may jointly consolidate gregarious behavior, requiring key signal transduction pathways and protein synthesis for memory consolidation (Geva et al., 2010; Ott et al., 2012), which is consistent with the view of plasticity as experience-dependent learning (Ellis, 1959).

Unpredictable environmental conditions and variable plant cover, typical of the desert locust’s arid habitat, are likely key drivers of the species’ extreme plasticity (Cisse et al., 2013, 2015; Sword et al., 2005). Unlike locust species common in grassland habitats, such as the Central American locust (*S. piceifrons*) and the migratory locust (*L. migratoria*), that tend to maintain cryptic solitarious phenotypes as long as conditions allow (Foquet et al., 2022; Guo et al., 2011), the desert locust’s response to crowding is notably faster than its response to isolation (Roessingh and Simpson, 1994). These transition dynamics help explain how desert locust aggregations remain stable when populations undergo frequent fission–fusion events under environmental constraints. Previous work has shown that even brief aggregation on resources can induce marked behavioral change in desert locusts, especially increased activity when food is scarce (Despland and Simpson, 2000b). In line with this, our data suggest that early differences (which in our case were most evident in group alignment rather than activity level or speed), became more pronounced over time and were likely reinforced through repeated interactions at feeding sites, where their association with food cues may provide synergistic value (Günzel et al., 2023; Petelski et al., 2024).

Large-scale transcriptomic studies have previously revealed broad differences between locust phases. In *S. gregaria*, nearly half of all transcripts are differentially expressed between solitarious and gregarious colonies (Bakkali and Martín-Blázquez, 2018). In *L. migratoria*, high-throughput sequencing across multiple tissues and larval stages narrowed phase-candidate genes to a smaller “PhaseCore” gene set (Yang et al., 2019). A recent cross-species analysis of these two major plague species found only a modest overlap in phase-related DEGs, though several enriched pathways and gene families were functionally shared (Bakkali et al., 2024). In contrast, our transcriptomics results identified a comparatively small set of differentially expressed genes. By rearing both groups in similarly enriched, semi-natural arenas that allowed foraging, roosting, and social encounters, we aimed to minimize confounding effects of confinement or isolation and to reveal behavioral and molecular differences directly linked to natural fluctuations in density and social experience. Future functional studies will be needed to clarify the role of these candidate genes in shaping behavioral phenotypes.

Out of the 157 DEGs we identified, only two genes overlapped with the phase-related DEGs common to both species reported in (Bakkali et al., 2024). Of these two, the Takeout-like protein (LOC126337065) is particularly noteworthy due to its juvenile hormone binding domain. In eusocial insects, juvenile hormone has been suggested as a regulator of caste differentiation, and members of the Takeout-like protein family are differentially expressed between castes (Ferreira et al., 2023). This protein family is diverse, shows varied expression across tissues, and exhibits complex regulatory interactions with JH. Our identified Takeout-like protein warrants detailed exploration. Existing literature offers several avenues through which Takeout-like proteins may induce or maintain phase change. Studies in *L. migratoria* have focused on the role of Takeout in chemosensation (Guo et al., 2011). Thus, it is plausible that the aggregation and local alignment behaviors we observed could be mediated by olfactory attraction, a sensory modality we found to be salient even in these otherwise visually dominant insects (Petelski et al., 2024). Beyond sensory modulation, Takeout may also play a role in metabolism and sleep-related activity, as shown in other species (Meunier et al., 2007; Sarov-Blat et al., 2000). This places Takeout at a central junction linking metabolic state and phase transition.

Reinforcing the proposed juvenile hormone–takeout axis for phase change, several other genes in the JH pathway also showed expression changes consistent with increased JH availability in the distributed condition. The major JH target, Krüppel homolog 1, also exhibited an upward trend in expression. In contrast to our findings, the involvement of JH in *S. gregaria* social plasticity remains debated (Breuer et al., 2003). Similarly, our dataset did not show clear evidence of a GABA–glutamate imbalance, and serotonin-related changes were limited. These discrepancies may reflect the transient nature of many neuromodulator shifts. Serotonin, for example, shows large spikes during gregarisation (thoracic ganglia) and solitarisation (brain), but stabilizes once phases are established (Rogers et al., 2004; Rogers and Ott, 2015). Similar dynamics have been reported for dopamine (Ma et al., 2011) and glutamate/GABA balance in L. migratoria (Yang et al., 2023). Such transient surges likely act as modulators of behavioral circuitry and as triggers for transcriptional programs, which can consolidate into the longer-term gene expression differences. Given the cellular heterogeneity of the nervous system, the granularity of single-cell RNA sequencing will be immensely valuable for future investigations to localize pathway engagement and resolve these discrepancies.

## 4 Materials and Methods

### 4.1 Animals

Desert locust nymphs *Schistocerca gregaria*(Forskål, 1775) obtained from a commercial breeder (B.T.B.E. Insektenzucht GmbH, Germany) were used in all experiments. Early 2nd instar locust nymphs were introduced into observation arenas (approximately 150 locusts per arena) in which they were kept for three weeks. Other than the arrangement of food and roosting stands, all extrenal conditions were kept similar between the arenas with a 12:12 h light:dark cycle, temperature of 27–29°C, and relative humidity of 45%. As a control with common gregarious and solitarious laboratory colonies, we kept additional 150 locusts from the same cohort in a gregarious holding container (50 × 50 × 80 cm cage) in our gregarious rearing facility, and 40 individuals in individual boxes (9 × 9 × 14 cm) in a separate, well-ventilated room with constant air exchange, as in (Petelski et al., 2024). All procedures were carried out in accordance with the “3Rs” principles, as stated in Directive 2010/63/EU and the German Animal Welfare Law.

### 4.2 Observation arenas, video recordings and motion capture system

Locusts were housed during the entire observation period in custom-built arenas (1.5 meters in diameter and 50 cm in height). To prevent the animals from climbing out, we applied petroleum jelly (Balea Vaseline, dm-drogerie markt GmbH, Germany) along the arena walls. Each arena was lit by two overhead lamps (NORKA Automation GmbH; continuous illumination, up to 8000 lux, 3000–6500 K), programmed for a 12L:12D cycle with a gradual ramp in intensity at the start and end of the light cycle, at 9:00 am and 9:00 pm.

The patches of food and roosting stands were arranged as follows: For the clumped condition, a single steel mesh structure (30 × 30 cm base size) and a food source were placed in the center of the arena. For the distributed condition, 17 small roosting meshes (10 × 12 cm each) were evenly spaced across the arena, each paired with a small food patch underneath (Fig 1A). The total roosting area and food weight were kept similar between both setups. Food was replaced daily under red light, and the arenas were cleaned every other day. Video recordings were made throughout the light period using a 12-megapixel camera (Iron 253 CoaXPress 12G 12MP rugged camera, Kaya Instruments, Haifa, Israel) with a 16-mm lens (IDS GmbH, Obersulm, Germany) at 5 FPS, operated using Streampix software (NorPix Inc, Montreal, H3K1G6 Canada). Furthermore, six motion capture cameras (Qualisys AB, 41105 Göteborg, Sweden) were mounted above the arenas to capture activity at 5 Hz, including during dark hours, through small reflective markers attached to the locusts during the final week of observation (days 16–23). The markers (3 mm in size) were cut from reflective polyester cloth and carefully attached to the locusts’ pronotum using removable gum. Marker count was checked daily, and individuals that lost their markers (mostly due to molting) were retagged. The cameras and motion capture system were calibrated before and after each experimental round.

### 4.3 Diet Composition and Preparation

Experiments were carried out with three different diets, selected to optimize tracking (trial 1, lab-made flat patches - T1), account for diet-related effects (trial 2, natural wheat pots - T2), and patch geometry (trial 3, cut wheat flat patches - T3). The laboratory-made diet (T1) contained a balanced mix of protein, carbohydrates, fiber, salts, vitamins, and water, following (Raubenheimer and Simpson, 2003; Simpson et al., 2002). Each batch was prepared using 50 g lentils (Bio rote Linsen, Edeka Baur, Germany), 30 g honey (Blütenhonig, Edeka Baur, Germany), 20 g wheat powder (Futterweizen, Pauls Muehle, Recklinghausen, Germany), 5 grams tamarind (TRS Tamarind, Thailand), 50 g gelatin (G2500, SIGMA-ALDRICH GmbH, Germany), and 1000 ml of water. The composition was selected based on initial screenings of locust development and survival rates. Lab-made diet food patches were prepared weekly, weighed, and stored at –4°C before being cut into 140 g portions, then placed in 15 cm (clumped) and 5 cm (distributed) Petri dishes, respectively. To assess possible diet-specific effects, Trial 2 and Trial 3 used natural wheatgrass: Trial 2:patches of wheatgrass grown in soil (625 cm^2^ and 49 cm^2^ for clumped and distributed), and Trial 3: Similar amounts of cut wheat in Petri dishes (same size as in T1).

### 4.4 Behavior assays

#### Individual preference assay

At the end of the observation period, individual behavioral phenotypes were assessed using a modified version of the assay adopted from (Roessingh et al., 1993b). Each locust was tested individually in a 100 cm-diameter circular arena with two patch choices: one containing 30 gregarious nymphs and one empty. The patches were placed equidistantly from the center, approximately 7 cm from the walls, with positions alternated between trials. Each trial lasted 15 minutes, during which the focal animal’s position was recorded by an overhead video camera, and after each reach trial, the arena was cleaned between trials.

#### Group Assay

Inter-individual interactions and group-level behavior were assessed towards the end of each trial in the absence of structured external cues. All roosting stands and food patches were removed, and the arenas were cleaned to remove any residue, while the locusts remained inside. The cleaning was performed early in the morning under red-light to minimize disturbance. Video recordings started with the beginning of the light cycle, and behavioral data were analyzed from 9:00 am in all trials.

### 4.5 Locust tracking - Trex and Qualisys Systems

Locust positions were tracked from the recorded videos using Trex (Walter and Couzin, 2021). Due to the large volume of video material, we selected two daily 40-minute segments for detailed analysis at the start of the light period and at peak activity in the afternoon. In Trial 1’s final observation week, we also used the Qualisys marker-based motion capture system, which provided high-resolution position data of the tagged locusts also during the dark hours. Data acquisition was conducted using Qualisys Track Manager (QTM) and further refined through a custom Unix shell script. To ensure reliability, we considered cross-validating the positional data obtained from TRex and Qualisys for consistency across systems. Tracked data from all experiments is available as CSV files on: https://github.com/Madhansai-Narisetty/Habitat_Phenotype_Data. Original video material is available upon request from the authors.

### 4.6 Data analysis

The preference Index (PI), estimating social preference in the individual assay was calculated as the proportion of time locusts spent on, or around, the box containing conspecifics minus the time at the empty box, divided by the sum of both. A range of one body length of 3cm around each patch edge was considered for the calculation. Preference values hence range from −1 to 1, with positive values indicating preference for the conspecifics box. Similar criteria was used for estimating time at the patches.

Tracking data from Trex and Qualisys systems were downsampled to 1 Hz for all analyses (except Fig 1, where data were downsampled to one every ten minutes to reduce autocorrelation). To capture robust behaviors and avoid artifacts from mistracking, locust instantaneous behaviors were detected by comparing Euclidean distances of position coordinates across consecutive frames as introduced in (Sayin et al., 2025). After filtering out ambiguous matches (e.g., when multiple individuals are closest to the same point), moving individuals were identified based on displacement thresholds (>0.5 cm/s and <10 cm/s). Global descriptions (heat maps, mean speed, and patch occupancy per frame) considered the entirety of the two arenas. Due to the strong thigmotaxis observed, the most outward section of the arenas was excluded from other analyses. The exclusion zone was determined based on the heat maps (Fig. 1 and 3), and the remaining total arena to be analyzed was 1.23 m^2^. Additionally, when movement analyses were considered in the presence of patches, locusts in proximity to patches were also excluded (patch radius + 3.75 cm for clumped, + 3.25 for distributed).

Alignment, defined here as circular collective movement with respect to the respective arena centers, were calculated according to (Buhl et al., 2006). To account for directional intermittency, absolute alignment values were reported here.

Scripts written in Python for behavioral analysis are available on GitHub LINK.

### 4.7 RNA Extraction and Sequencing

Whole brains were extracted at the end of Trial 1 and 3. For the extraction, 80 animals from each condition were sampled (pooled into biological replicates of 10 brains each; yielding 8 replicates per condition). Brains were rapidly dissected in ice-cold DEPC PBS (made with MilliQ water), and placed in 100*µ*L TRIzol (Thermo Fisher Scientific), homogenized immediately using a mechanical pestle and stored at −80 degrees. RNA extraction was performed using the Purelink DNase and RNA purification kits (Thermofisher Scientific) according to kit protocols. RNA samples were shipped to Novogene GmbH (Munich, Germany) for mRNA sequencing (Illumina Novoseq X Plus platform, 30 mio PE150 read pairs). RNA-Library preparation with Poly-A mRNA enrichment tail, library quality control, read assembly was performed by Novogene. One sample was excluded due to very low mapping yield (Fig. S1B).

Differential gene expression was performed with Deseq2, accounting for batch effects, (Love et al., 2014), after “ribosomal RNA” hits were filtered out. Enriched pathways (GSEA) were explored using with KEGG pathway data for (clusterProfiler (Yu et al., 2012), KEGGREST (Tenenbaum, 2025) packages). A ranked gene list from DESeq2 results was analyzed without an adjusted cutoff to inspect all nominal hits, and the top 20 pathways were selected based on the raw gseKEGG p-value. Counts, here as in overlaps with significantly differentially expressed genes, were quantified using a nominal gene-level threshold of p < 0.1 determined by DESeq2. Individual enriched pathways were then visualized using RDkit (Landrum et al., 2006) in Phyton based on *Schistocerca gregaria* KEGG pathways (Kanehisa and Goto, 2000).

We compared DEGs shared between this study and a list of common phase-dependent DEGs found in both *Schistocerca gregaria* and *Locusta migratoria*. After extracting FASTA sequences from (Bakkali et al., 2024), *S. gregaria* contigs were matched to NCBI reference protein (or RNA) sequences with blastx (or blastn for anonymous genes; see the reference aforementioned for details) based on the first best hit. The resulting gene IDs are then matched with our DESeq2 results.

## 5 Acknowledgments

We wish to thank Mathias Gunther, Heike Naumann, Nina Schwarz, Hannes kübler, Elvira Mamanova, Jayme Weglarski, Dominique Leo, Alexander Bruttel, and Cyrill Fromm for their assistance in the project. Jackob Davidson and Angela Albi for their insight on Tracking. Anna Stockl, Iain Couzin and Greg Sword for insightful suggestions.

---

**Figure S1:**
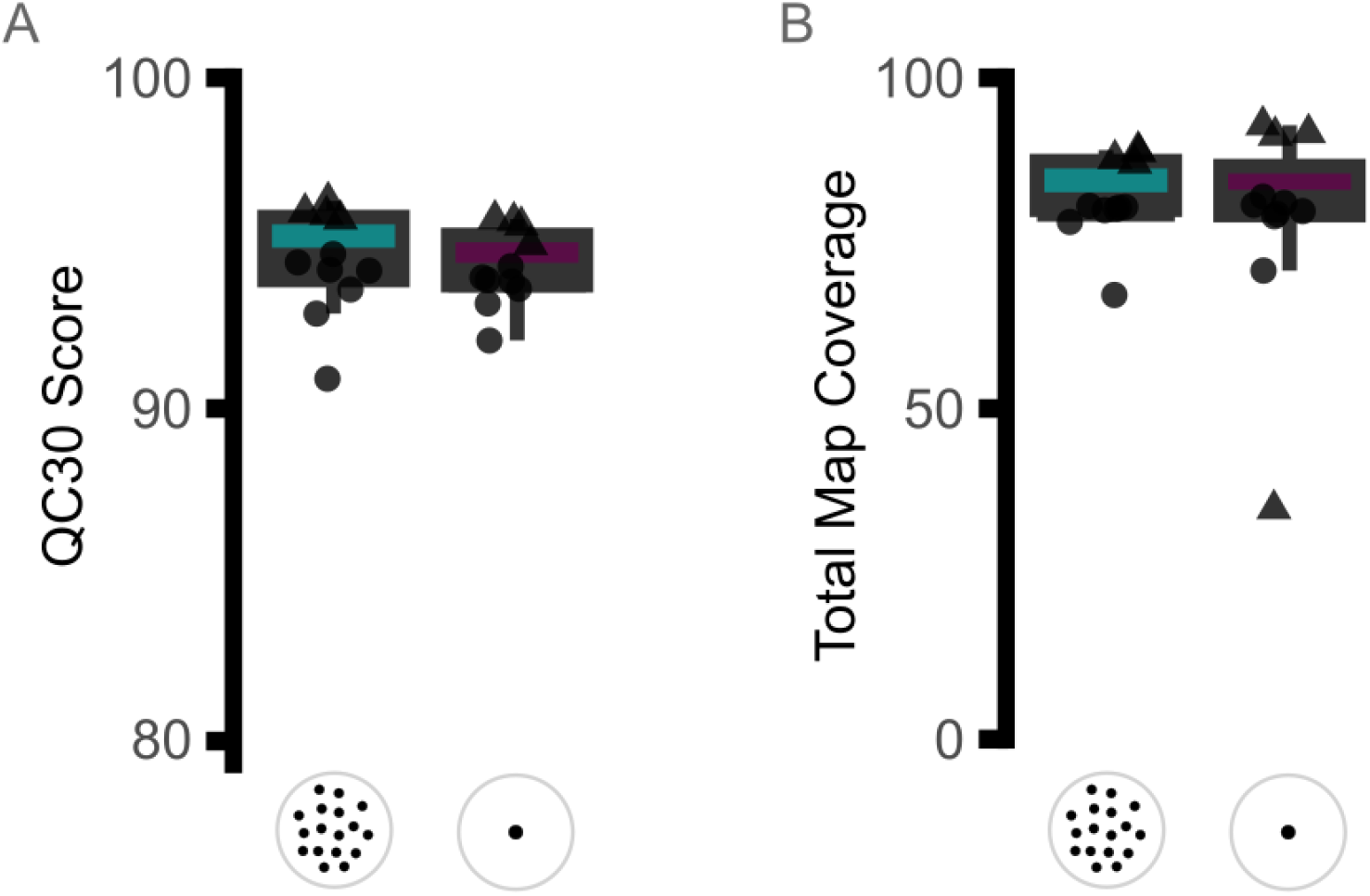
Quality Assessment of RNA Sequencing. (A) Q30 score. The proportion of reads with a sequencing accuracy greater than 99.9 per cent. (B) Total mapping coverage. One sample from the second batch was excluded from the study due to extremely low mapping to the NCBI reference genome. Each point represents an individual sample within its respective trial.

**Figure S2:**
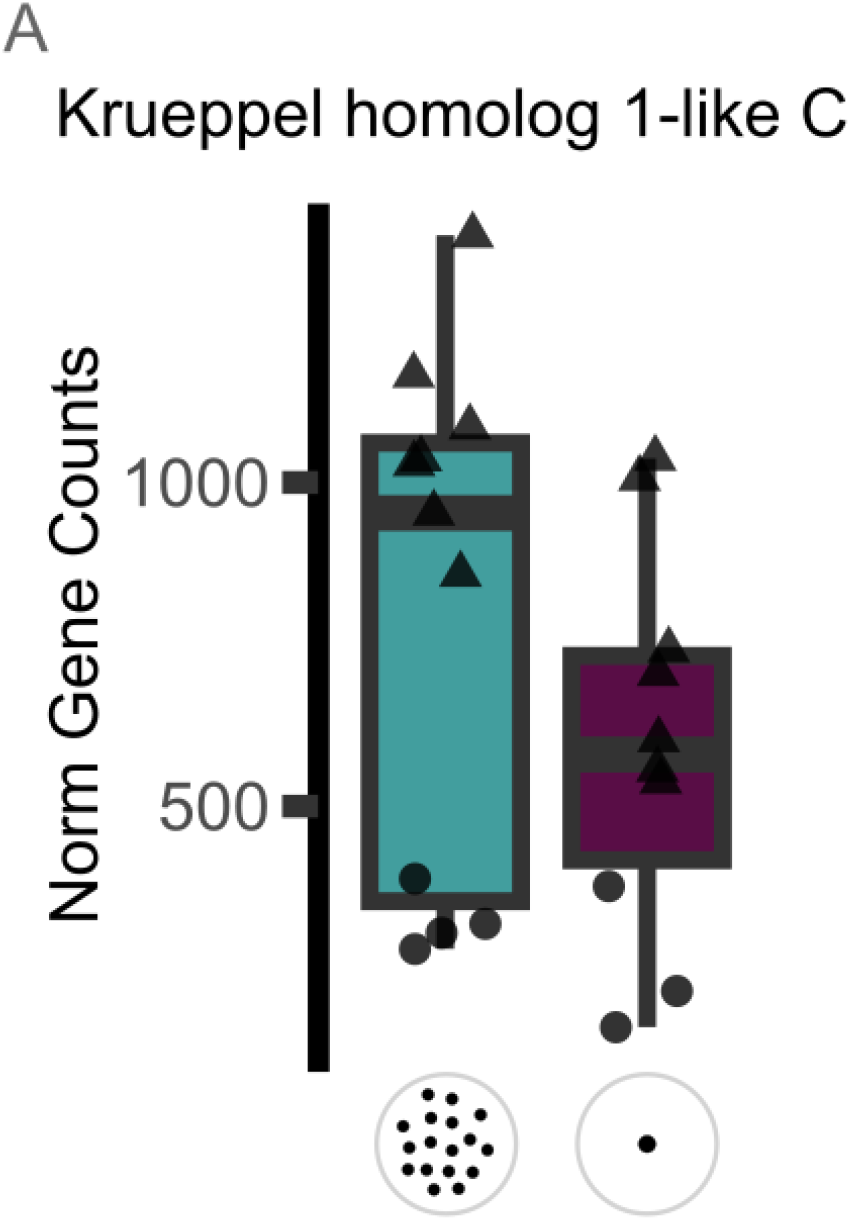
Normalized counts for Krueppel homolog 1. 2. (A) Juvenile hormone target Krueppel homolog 1 (LOC126272608) was upregulated in distributed condition (log_2_ FoldChange > −0.4986, adjusted *p*-value <0.0549). Each point represents an individual sample within its respective trial.

**Figure S3:**
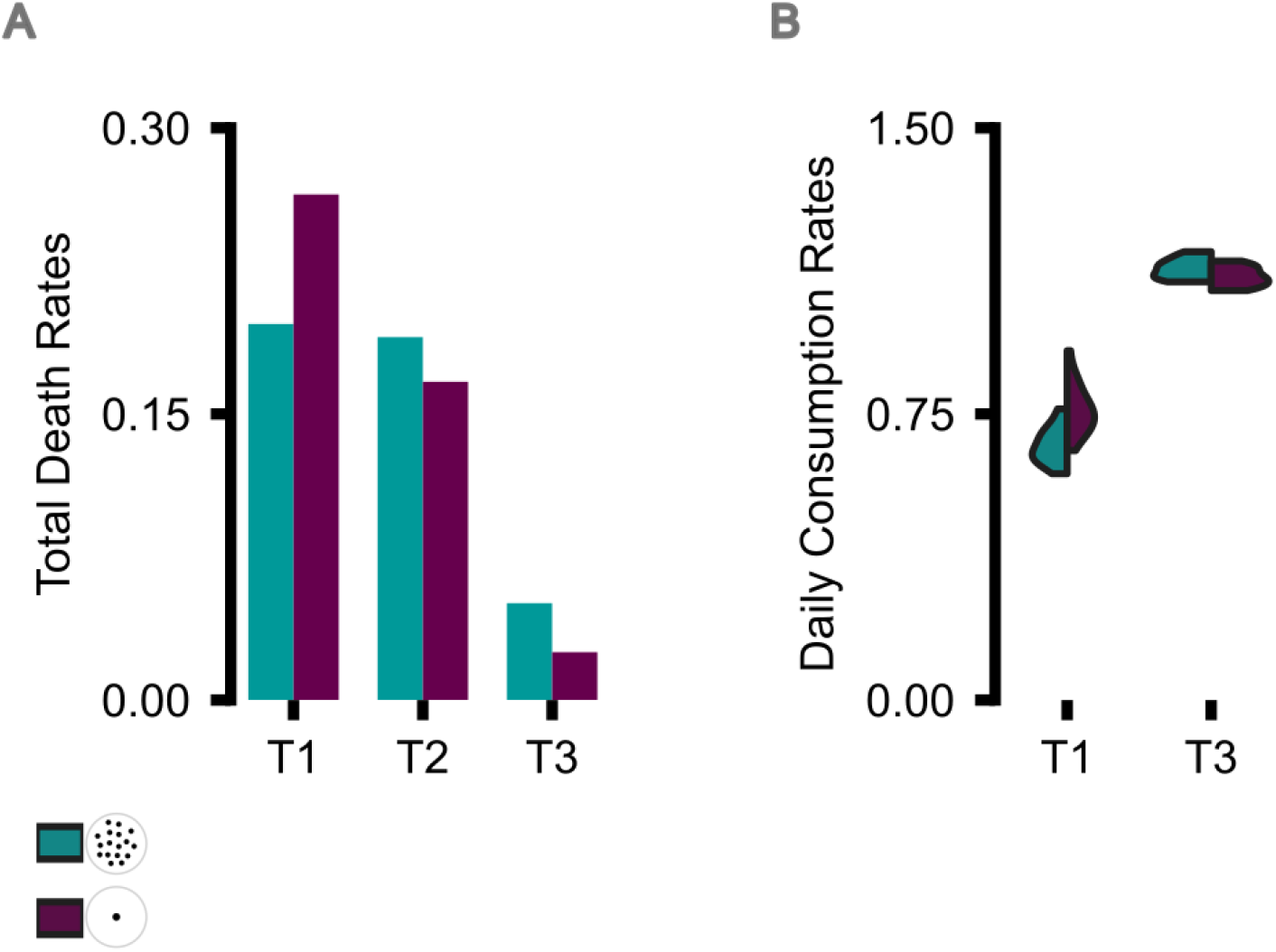
(A) Mortality rates in all trials (B) Average rate of total daily food consumption for each arena in T1 and T3.

